# Mathematical model of replication-mutation dynamics in coronaviruses

**DOI:** 10.1101/2024.01.29.577716

**Authors:** K.B. Blyuss, Y.N. Kyrychko

## Abstract

RNA viruses are known for their fascinating evolutionary dynamics, characterised by high mutation rates, fast replication, and ability to form quasispecies - clouds of genetically related mutants. Fast replication in RNA viruses is achieved by a very fast but error-prone RNA-dependent RNA polymerase (RdRP). High mutation rates are a double-edged sword: they provide RNA viruses with a mechanism of fast adaptation to a changing environment or host immune system, but at the same time they pose risk to virus survivability in terms of virus mutating beyond its error threshold. Coronaviruses, being a subset of RNA viruses, are unique in having a special enzyme, exoribonuclease (ExoN), responsible for proofreading and correcting errors induced by the RdRP. In this paper we consider replication dynamics of coronaviruses with account for mutations that can be neutral, deleterious or lethal, as well as ExoN. Special attention is paid to different virus replication modes that are known to be crucial for controlling the dynamics of virus populations. We analyse extinction, mutant-only and quasispecies steady states, and study their stability in terms of different parameters, identifying regimes of error catastrophe and lethal mutagenesis. With coronaviruses being responsible for some of the largest pandemics in the last twenty years, we also model the effects of antiviral treatment with various replication inhibitors and mutagenic drugs.

## 1. Introduction

Among various known viruses known to infect humans, a particular place is occupied by coronaviruses (CoV), which are enveloped positive-sense RNA viruses of the *Nidovirales* order, with genome sizes of 26-32 kB, which makes them some the largest and genomically most complex RNA viruses [69]. At present, seven different coronaviruses have been identified, four of which, namely, HCoV-229E, -OC43, -NL63, and -HKU1 are known to cause common cold. The other three are SARS-CoV that caused an epidemic in 2003, MERS-CoV identified in 2012, and SARS-CoV-2 first identified in late 2019 that has caused the COVID-19 pandemic [55, 49]. The life cycle of coronaviruses proceeds as follows. Upon cell entry through either endosomes or membrane fusion, genomic viral RNA is uncoated, and open reading frames ORF1a and ORF1b are immediately translated into polyproteins pp1a and pp1b that are then cleaved into individual non-structural proteins (nsps) [106, 33]. Some of these nsps form the so-called replication-transcription complex (RTC), in which viral RNA is replicated and transcribed into sub-genomic RNAs that then form part of the virions [60, 20]. Importantly, since host cells lack the machinery to translate viral mRNA into proteins, all RNA viruses encode their own RNA-dependent RNA polymerase (RdRp). In CoV, RdRp is encoded by nsp-12, and once RTC has been formed, this RdRp uses available (+)-strand of viral RNA as a template to produce a negative-sense strand, which, in turn, is then used for further production of a positive-sense strand, and this is how viral replication is achieved [69, 76, 42].

A defining feature of RNA viruses in general, and CoV in particular, is their extremely high speed of replication, which is accompanied by high rates of mutation [22, 23, 26, 24] that are underpinned by the lack of proofreading ability of RdRp [27, 82, 83]. This means that during replication, when forming new sense and anti-sense RNA strands, RdRp can erroneously attach incorrect nucleotides to the growing strands. The result of high rates of mutations is the formation of so-called *molecular quasispecies* [28, 29], a term used to denote a cloud of genetically related mutants. Heterogeneous structure of quasispecies provides a pool of phenotypes able to adjust to environmental change and selection pressure from host. While being seemingly an effective survival strategy, mutations cannot increase without bounds due to the existence of *error threshold* [44, 26], beyond which further selection is impossible, as mutant genomes would outcompete mutation-free genomes. Once mutation rates exceed those at the error threshold, this results in extinction of the highest-fitness genotype, known as an *error catastrophe* [29, 26], while genotypes with lower fitness that have a higher robustness to mutations can still be maintained. Bull et al. [5] provide a nice discussion of error threshold, error catastrophe and extinction threshold. Solé et al. [96] have studied a model with master and mutant strands that have different replication rates and showed a transition from quasispecies to the state, where only mutant population is present, depending on the relation between mutation rate and the fitness rate of the master strain. Error catastrophe has been also analysed in the literature as a phase transition, using methodology from statistical physics [2, 97, 98].

RNA viruses are known to operate close to the error threshold [44, 26, 24, 14], which is both an advantage, and a potential vulnerability for them. An important concept in this context is that of *lethal mutagenesis* that denotes loss of viral infectivity [56], usually as a result of chemically-induced mutagenesis. While this resembles error catastrophe, the distinction between these two concepts is that error catastrophe rather describes a qualitative reorganisation of genotype space as a result of evolution, where extinction of particular genotypes may or may not happen depending on their fitness and mutation rates, whereas lethal mutagenesis is specifically concerned with driving population to extinction as a result of mutations [6]. Lethal mutagenesis arising from error catastrophe has been proposed as one possible antiviral strategy that can result in extinction of RNA virus [25, 94]. Experimentally, it has been shown how chemically-induced lethal mutagenesis can extinguish a number or different RNA viruses, including hepatitis C virus [14], HIV [56], foot-and-mouth disease virus (FMDV)[91, 70], and lymphocytic choriomeningitis virus [41, 79].

Unlike any other RNA viruses, CoV are quite unique in having Exoribonuclease (ExoN), an enzyme formed by non-structural proteins nsp-10 and nsp-14 encoded in the viral RNA [50, 69, 66, 93]. While, coronavirus RdRps are extremely fast due to inherent lacking of fidelity, ExoN is able to significantly improve the fidelity of viral replication by removing (excising) incorrect nucleotides attached to the growing RNA strands by error-prone RdRp. RdRp encoded by nsp-12 of SARS-CoV is the fastest known RdRp among any RNA viruses, but its error rate, i.e. the percentage of incorrect nucleotides being attached to the nascent RNA strand, exceeds equivalent error rates for other viral RdRps by an order of magnitude [90]. Hence, for this virus to survive, it is essential that RdRp-induced replication errors are corrected by the ExoN. In essence, replication of CoV is an ongoing ‘tug-of-war’ between mutations produced by RdRp, and proofreading and error correction performed by the ExoN.

A number of mathematical models have studied full intracellular life cycle of RNA viruses, including hepatitis C (HCV), dengue (DENV) and SARS-CoV-2 as examples of (+)-sense RNA viruses, and influenza A as an example of (-)-sense RNA viruses. One of the earliest such models is the model of Dahari et al. [15], which considered the dynamics of HCV replication in Huh7 cells, which included various stages of virus life cycle, including intra-cytoplasmic translation, formation of RTCs, individual production of positive and negative RNA strands, as well as structural proteins. Binder et al [4] extended this model to explore an important role of the timescales associated with processing transfected RNA and forming RTC. Zitzmann et al. have used a version of the same model to analyse the effects of exosomal and viral RNA secretion in HCV [117], as well as to study the within-cell dynamics of dengue virus replication, with account for host immune responses [116]. Very recently, Zitzmann et al. [118] considered a generic model of intracellular viral replication and showed that with very minor virus-specific changes, the model could be fit quite well to the measurements of viral replication of HCV, DENV and Coxsackie virus B3 (CVB3). This model has provided important insights into shutting down mRNA translation in host cells being a vital factor in controlling the efficiency of viral replication. Grebennikov et al. [42] have analysed a similar kind of model for SARS-CoV-2 and fitted parameters to reproduce viral kinetics observed *in vitro* in experiments in MHV- and Vero E6-infected cells.

During intracellular replication, the same viral RNA is involved in three different, mutually exclusive, processes, namely, translation into proteins, being used as a template for further replication, and being packaged into new virions. Hence, an important question is how the available viral RNA can be allocated and possibly dynamically re-allocated to each of these three activities to ensure optimal performance in terms of enhancing the production of successful new virus particles. McLean et al. [65] have explored in details such trade-off in RNA allocation using a simplified version of Dahari’s model for HCV. Nakabayashi [68] used an example of HCV to study the same problem of how viral RNA can be distributed to those three functions involved in viral replication. He considered the possibilities of explosive and arrested replication, showing how viral RNA should be re-distributed to avoid replication arrest.

While providing important insights into various stages of viral replication, all of the above models only considered the actual process of viral replication from an RNA template but did not include the possibility of mutations, which, with low-fidelity RdRp of RNA viruses, may play a significant role in viral dynamics. To investigate this specific issue, Sardanyés et al. [84] proposed and analysed a mathematical model of viral replication within RTC for positive-sense single-stranded RNA viruses, with account for mutations that can occur on either of two strands. That work also noted that when analysing viral replication with mutations, it is essential to accurately represent so-called *viral replication modes* that characterise which template is used to generate viral progeny [84]. Two extreme cases of viral replication modes are the Stamping Machine Replication (SMR) and the Geometric Replication. In the case of SMR, first proposed by Stent [99], positive-sense viral RNA that comes from the viral particle after uncoating is transcribed into complementary negative strands, and it is only these negative strands that are then used as templates for subsequent production of the entire viral progeny. In contrast, GR as proposed by Luria [58] describes a situation where both sense and anti-sense strands of viral RNA are used as templates for further rounds of viral replication within the RTC. Replication mode of some RNA viruses, such as vesicular stomatitis virus [12] and the poliovirus [88] is quite close to GR, with multiple rounds of RNA copying inside the cell. Other RNA viruses, such as bacteriophages Q*β* [36] and *ϕ*6 [8], as well as the turnip mosaic virus [64], replicate in a way that is very reminiscent of SMR. At the same time, since many other viruses operate in a regime somewhere between those two extremes, hence, it is convenient to represent the mode of replication using a continuous parameter 0 *< α ≤* 1 [85], with *α* ⪆ 0 corresponding to SMR, and 0 *< α ≤* 1 describing GR, with *α* = 1 being a situation, where both positive and negative strands replicate at the same rate.

Several papers have considered replication-mutation dynamics using the model of Sardanyés et al. [84] and its modifications to explore various scenarios of replication dynamics depending on the viral replication mode [85, 87, 35, 105]. A common feature of these models is the extinction of viral genomes through a non-degenerate transcritical bifurcation that separates an extinction steady state from a stable steady state, in which both master and mutant strands are simultaneously present [85]. Since this happens as a smooth transition between stable steady states under a change in parameter values, this suggests a presence of a second-order phase transition. Similar behaviour has been observed in other quasispecies models of RNA viruses with mutations [102, 80]. An important ingredient in models of virus replication-mutation dynamics is that of *fitness landscape*, which characterises the distribution of viral fitness of mutant strands, as quantified by their replication rate in relation to that for the master strand. The majority of the earlier-mentioned papers have used the Swetina-Schuster single-peak fitness landscape [101], in which mutants with different numbers of mutations are all grouped together, and they all have the same replication rate that is lower than or equal to that of the master strand, irrespective of the number of mutations in a particular mutant. Naturally, such an assumption is a simplification and one could consider more complex, time-varying or rugged landscapes [53]. However,

Swetina-Schuster fitness landscape has been successfully used to study quasispecies dynamics in RNA viruses [5, 71, 96]. Sardanyés et al. [84] have investigated an interplay between replication modes and fitness landscapes, which were characterised by the fitness expressed as a linearly decreasing function of the Hamming distance between the mutant and master genomes when represented as binary strings. Thébaud et al. [105] have explored an inter-connection between optimal replication strategy and the rates of mutations in positive-sense single-stranded RNA viruses.

In this paper, we focus on mathematical modelling of replication dynamics of CoV, with account for possible mutations during replication. Our model is based on an earlier work of Fornés et al. [35], but it also specifically considers the role played by ExoN, a feature that is unique to coronaviruses and the one that provides them with a mechanism of repair for incorrect nucleotides being attached to nascent RNS strands. Using Swetina-Schuster single-peak fitness landscape will allow us to group a cloud of different mutants into an aggregate mutant sequence having a fitness that is lower than or equal to that of the master sequence. We will study different types of mutations, and in each case we will analyse steady states of the model and their stability. We will also investigate the effects of drugs that have been or potentially can be used to treat disease caused by CoV. In the next section, we derive the model

## 2. Mathematical model

Focusing on CoV replication inside the RTC, we follow Fornés et al. [35] and consider overall viral population to be represented by four classes of RNA: master and mutant positive-strand classes, and master and mutant negative-strand classes. Polarity of a given RNA strand will be denoted by *p* for positive strands, and by *n* for negative strands, while subindex 0 or 1 will indicate, whether we are dealing with a master strand, or a mutant strand. With these notations, *p*_0_ and *n*_0_ denote concentrations of positive- and negative-sense single-stranded RNA with the master sequence, and *p*_1_ and *n*_1_ denote corresponding concentrations for the mutant sequence. We take *k*_0_ *>* 0 and *k*_1_*≥* 0 to be replication rates of the master and mutant sequences, respectively, and the mutation rate is denoted as *µ*. Mutations can be classified as: *neutral* (*k*_0_ = *k*_1_ = 1), which do not cause a reduction in replication rate for the mutant strand, *deleterious* (0;≲ *k*_1_ *< k*_0_ = 1), where mutants have a lower replication rate, and *lethal* (*k*_1_ = 0), where mutant strands cannot replicate. With negative strand being replicated from the positive strand, and the positive strand being replicated from the negative strand, the parameter 0 *≤α ≤* 1, mentioned in the introduction, is used to denote the ratio of two replication rates, with the strands replicating at the same rate for *α* = 1, which is known as geometric replication (GR), and when 0;≲ *α ≪*1, there would be a much stronger replication of the positive strand from the negative strand than the other way around, known as a stamping machine replication (SMR).

Fornés et al. [35], as well as other works based on Sardanyés et al. [84], only considered forward mutations, i.e. mutations from the master strand into the mutant strand. This assumption was based on the argument that with the length of RNA viral genome being of the order of 10^6^ nucleotides, the probability of backward mutations would be extremely low. A number of other models that studied the dynamics of quasispecies used the same assumption [5, 71, 96]. In our case, while we will also not directly include mutations from the mutant strands into the master strands of complementary polarity, we will allow for the transition from mutant to master strands of the same polarity under the action of ExoN.

If we denote by [*NSP*] the total amount of non-structural proteins in the RTC and assume that for the duration of replication-transcription cycle the concentration of nsps in the cell is constant, we then denote by [*RdRp*] = *σ*_1_[*NSP*] and [*ExoN*] = *σ*_2_[*NSP*] the proportions of RdRp and ExoN enzymes, respectively. Assuming a saturated (Michaelis-Menten) kinetics, as is common in enzyme-catalysed reactions [13, 51], the effect of RdRp and ExoN on strand replication and transformation from mutant into master strand, can be written as

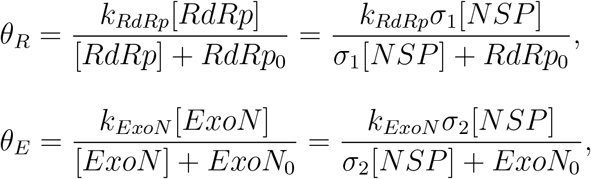

where *k*_*RdRp*_ and *k*_*ExoN*_ are kinetic constants, and *RdRp*_0_ and *ExoN*_0_ are half-maximal concentrations of RdRp and Exon, respectively.

The model can now be written in the form

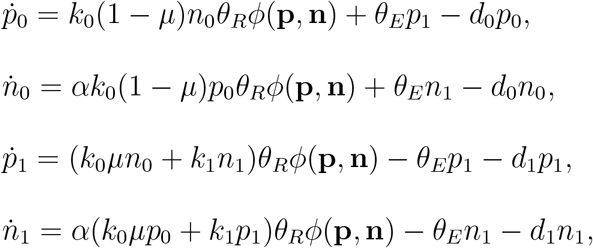

where *d*_0_ and *d*_1_ are natural degradation rates for master and mutant strands, and the factor

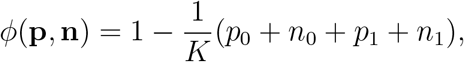

is introduced to describe the competition between viral strands for cell proteins, and to bound the growth. We can rescale all strand variables with the carrying capacity *K*, or simply assume *K* = 1. For simplicity, we will also assume degradation rates of master and mutant strands to be the same, i.e. *d*_0_ = *d*_1_ = *d*, while the replication rate of the master strand is assumed to be equal to one, *k*_0_ = 1. Rescaling the time with 1*/d*, introducing rescaled parameters 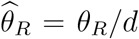 and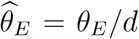, and dropping the hats for notational convenience, we then have a modified model

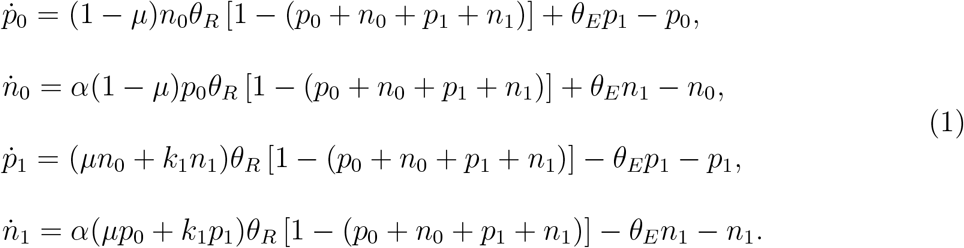

This system is well-posed, with its solutions being non-negative and bounded for non-negative initial conditions. The difference between this model and the model considered by Fornés et al. [35] lies in the explicit inclusion of *θ*_*R*_ to represent the role of RdRp in viral replication, and in the presence of terms with *θ*_*E*_ as a factor, to account for ExoN-induced transitions from mutant strands back to master strands.

## 3. Steady states and their stability

For any parameter values, the system (1) has an extinction steady state *E*_0_ = (0, 0, 0, 0), where viral strands with both master and mutant sequence go extinct.

Jacobian of linearisation near any of the steady states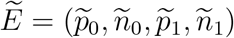 has the form

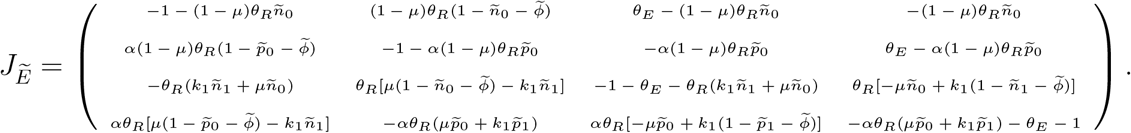

For the extinction steady state *E*_0_, the characteristic equation is

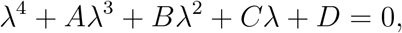

where

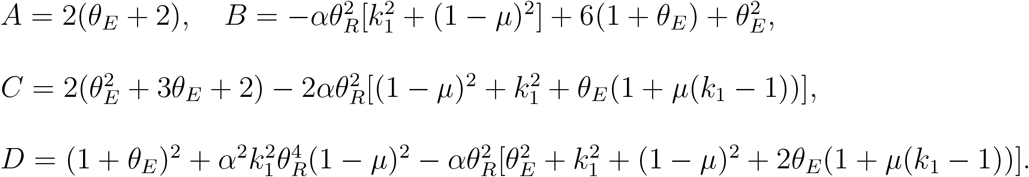

Conditions for stability of the steady state *E*_0_ can now be found from the Routh-Hurwitz criteria:

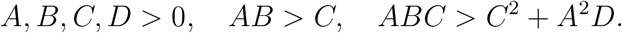

The last two conditions have the explicit form

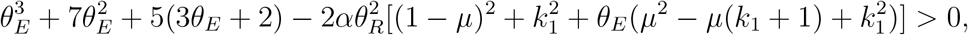

and

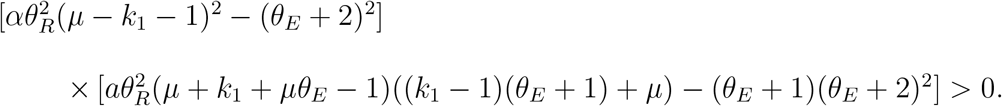

In the case when *θ*_*E*_ = 0, i.e. ExoN is not effective at proofreading and turning mutant strands back into the master strands, there is also a possibility of a steady state where master strand goes extinct, 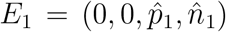, which is similar to the case considered by Fornés et al. [35]. The mutant-only steady state *E*_1_ has components

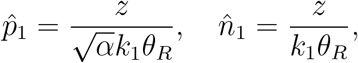

where

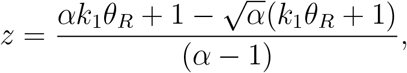

and is biologically feasible for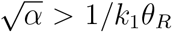. Another possibility is the mixed steady state 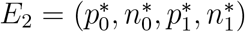, in which all components are greater than zero.

Stability conditions for the extinction steady state *E*_0_ in the case of vanishing *θ*_*E*_, which biologically corresponds to the situation where ExoN is either blocked or ineffective at correcting replication errors caused by RdRp, reduce to a single condition

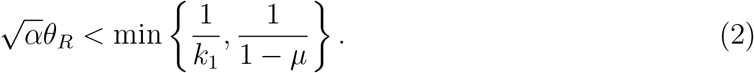

Obviously, this condition can only be satisfied, provided *θ*_*R*_ *>* 1, and otherwise, the extinction steady state *E*_0_ will be stable for all possible values of *α, k*_1_ and *µ* in the case where ExoN is either absent or non-functional, i.e. for *θ*_*E*_ = 0. We note that the loss of stability of *E*_0_ through crossing the boundary 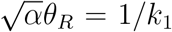 coincides with the mutant-only steady state *E*_1_ becoming biologically feasible. Stability condition for the mutant-only steady state *E*_1_ is given by

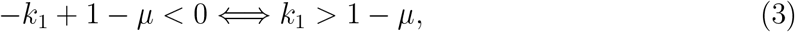

as the remaining characteristic eigenvalues can be shown to be negative for any values of the parameters.

Figure 1 illustrates transitions observed for *θ*_*E*_ = 0 between a stable mixed steady state *E*_2_, a steady state state *E*_1_, in which the master strand has gone extinct, and only the mutant is present, and the steady state *E*_0_, corresponding to extinction of both master and mutant sequences. For sufficiently small mutation rates *µ*, we observe that for very small values of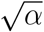, corresponding to SMR regime, the system exhibits extinction of all genotypes regardless of whether mutations are neutral or deleterious. An intuitive explanation for this is that the negative strands are not produced quickly enough to ensure subsequent replication of positive strands before all strands are degraded. For higher values of 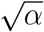, once the value of 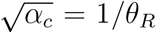 is crossed, we observe the birth of the mutant-only steady state, which exists for values of *k*_1_ close to 1 (almost neutral mutations). For sufficiently high values of 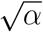 exceeding the threshold value of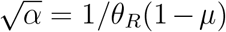, i.e. moving to the GR regime, there is a stable mixed steady state *E*_2_, where both master, and mutant sequences are present. In the parameter region, where *E*_1_ is also feasible, the steady state *E*_2_ coexists with an unstable *E*_1_ for more deleterious mutations (lower *k*_1_). There are two boundaries, through which the mixed steady state *E*_2_, describing viral QS, can disappear: as mutations become less deleterious, and hence, the fitness of mutants *k*_1_ increases, eventually, *E*_2_ will cross the error threshold at *k*_1_*c* = 1 *− µ*, resulting in the mutant-only steady state *E*_1_. Another possibility fordisappearance of *E*_2_ is when the replication rate 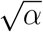 is reduced below the critical value of 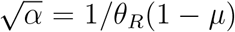, at which point we observe lethal mutagenesis, as represented by the stable extinction steady state *E*_0_. We note that provided replication rate *α* is sufficiently high, the viral QS can be maintained (in the case of not too high mutation rates) even for almost lethal and lethal mutations that result in mutants having effectively zero fitness. Increasing the mutation rate *µ* reduces the range of parameters, where viral QS can be maintained, and once the value of *µ* exceeds the threshold value of 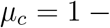 1*/θ*_*R*_, viral QS as represented by the mixed steady state *E*_2_, does not exist for any mutant fitness *k*_1_ and any replication rate *α*. In that case, the system can exhibits either an extinction steady state *E*_0_ for values of *k*_1_ and *α* below the boundary 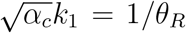, and the mutant-only steady state *E*_1_ above this boundary.

**Figure 1.**
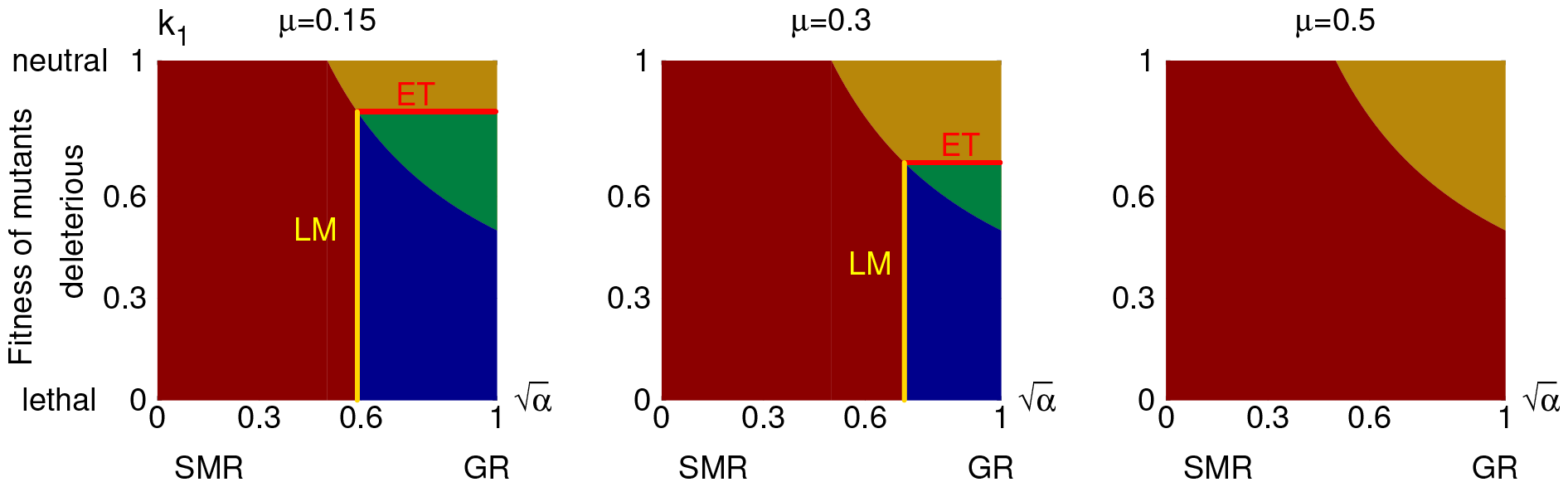
Regions of stability of different steady states of the model with *θ*_*E*_ = 0 and *θ*_*R*_ = 2. Colours indicate a stable extinction steady state *E*_0_ (dark red), stable mutant-only steady state *E*_1_ (golden), stable mixed steady state *E*_2_ with unstable *E*_1_ (green), and stable mixed steady state *E*_2_ with infeasible *E*_1_ (blue). Lines indicate an error threshold (ET) in red, and the transition from a stable mixed steady state *E*_2_ to a stable extinction steady state *E*_0_ through lethal mutagenesis (LM). Modes of replication are denoted as SMR (stamping machine replication) and GR (geometric replication).

In Fig. 2 we have explored the same case of *θ*_*E*_ = 0, but focused instead on copying fidelity, as given by (1 *− µ*), and the replication rate *α* for different value of mutant fitness *k*_1_. Similarly to the previous figure, in the case of SMR, described by low replication rates *α*, the only possible outcome for the system is the extinction of both master and mutant sequences. For values of *k*_1_ lower than the 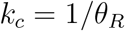, including the case of lethal mutations *k*_1_ = 0, the system only exhibits either an extinction steady state *E*_0_ for values of copying fidelity (1 *− µ*) and replication rate 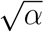 lower than the boundary 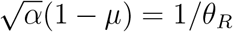, or the mixed steady state *E*_2_ above this boundary. The boundary itself represents the curve of lethal mutagenesis, associated with a transition from the viral QS to complete extinction. For values of *k*_1_ higher than *k*_*c*_, there is an interval of replication rates, close to the GR case, where for high copying fidelity one has stable viral QS, while for lower copying fidelity, there is a stable mutant-only steady state *E*_1_. An explanation of this situation lies in observing that very high replication rates *α* ensure the existence of viral sequences, but if the fidelity of replication is not sufficiently high (or, alternatively, mutation rate *µ* is high enough), the system is able to replicate mutant sequence but cannot maintain the master sequence, thus resulting in a stable mutant-only steady state. The transition from lower fidelity to higher fidelity through the boundary 1 *− µ*_*c*_ = *k*_1_ corresponds, therefore, to crossing an error threshold. As the mutant fitness *k*_1_ increases, the range of replication rates *α*, for which one observes the stable mixed steady state and its transition into a mutant-only steady state for lower copying fidelity, grows. Eventually, in the case of neutral mutations, *k*_1_ = 1, the mixed steady state completely disappears, and there is only a stable extinction steady state *E*_0_ for replication rates lower than 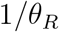 and a stable mutant-only steady state *E*_1_ for values of replication rates higher than 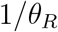.

**Figure 2.**
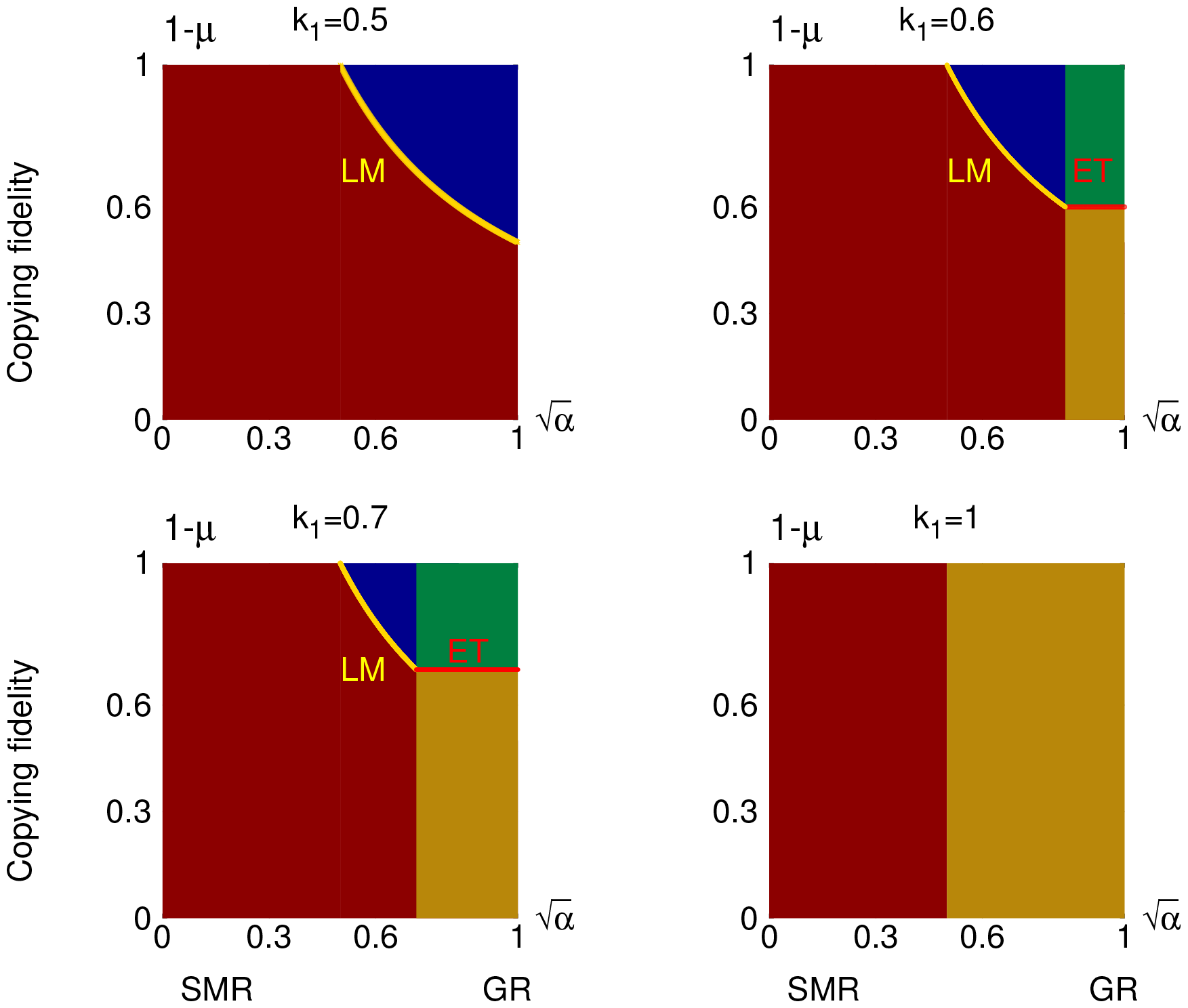
Regions of stability of different steady states of the model with *θ*_*E*_ = 0 and *θ*_*R*_ = 2. Colours indicate a stable extinction steady state *E*_0_ (dark red), stable mutant-only steady state *E*_1_ (golden), stable mixed steady state *E*_2_ with unstable *E*_1_ (green), and stable mixed steady state *E*_2_ with infeasible *E*_1_ (blue). Lines indicate an error threshold (ET) in red, and the transition from a stable mixed steady state *E*_2_ to a stable extinction steady state *E*_0_ through lethal mutagenesis (LM). Modes of replication are denoted as SMR (stamping machine replication) and GR (geometric replication).

For *θ*_*E*_ *>* 0, the mutant-only steady state *E*_1_ would not be feasible, because under the action of ExoN, master sequence strands would always be recovered. In this case, the only possible non-trivial steady state is the mixed equilibrium 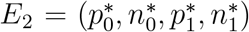. In the case of lethal mutations, characterised by *k*_1_ = 0, the components of the mixed equilibrium can be found explicitly as

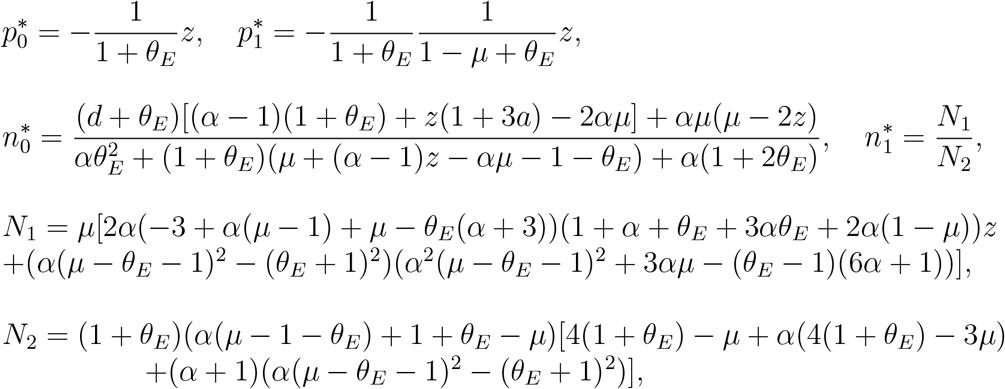

where

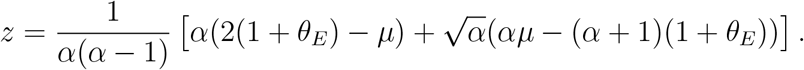

For *θ*_*E*_ *>* 0 and non-lethal mutations (*k*_1_ *>* 0), it does not prove possible to find a closed form expression for the mixed steady state *E*_2_, though extensive numerical simulations suggest that in all parameter regions we explored, the mixed steady state *E*_2_ was unique when it was feasible.

Figure 3 illustrates transitions between mixed and extinction steady states in the case when ExoN is present and functional, i.e. for *θ*_*E*_ *>* 0. We observe that as the replication rate *α* increases, the lethal mutagenesis occurs for smaller values of *θ*_*R*_, in agreement with condition (2), i.e. a larger part of the parameter plane is occupied by the stable mixed steady state *E*_2_. As *α →* 1, the value of *θ*_*R*_, at which the transition from *E*_2_ to *E*_0_ is taking place for *θ*_*E*_ *→* 0, tends to the smaller of 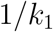 and 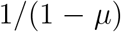. Plot 3(a) suggests that for very small but positive values of *θ*_*E*_, in other words, as ExoN only starts to be effective at correcting nucleotide errors caused by the RdRp, lethal mutagenesis occurs for a slightly smaller value of *θ*_*R*_ than when *θ*_*E*_ = 0. However, there appears to be no further variation in the boundary of lethal mutagenesis for higher values of *θ*_*E*_. When considered in the parameter plane of copying fidelity (1 *− µ*) and 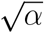, the extinction steady state occurs for smaller values of 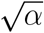 and for lower copying fidelity. This is to be expected, as effectively it means that replication fidelity is not sufficient to maintain effective replication of both master and mutant sequences, and with the mutant-only state being unavailable due to the action of ExoN that would result in the recovery of the master sequence, the only other possibility is the extinction of both sequences. Increasing *θ*_*E*_, i.e. making ExoN more effective at correcting RdRp-induced replication errors, allows the mixed steady state *E*_2_ to persist for lower values of copying fidelity.

**Figure 3.**
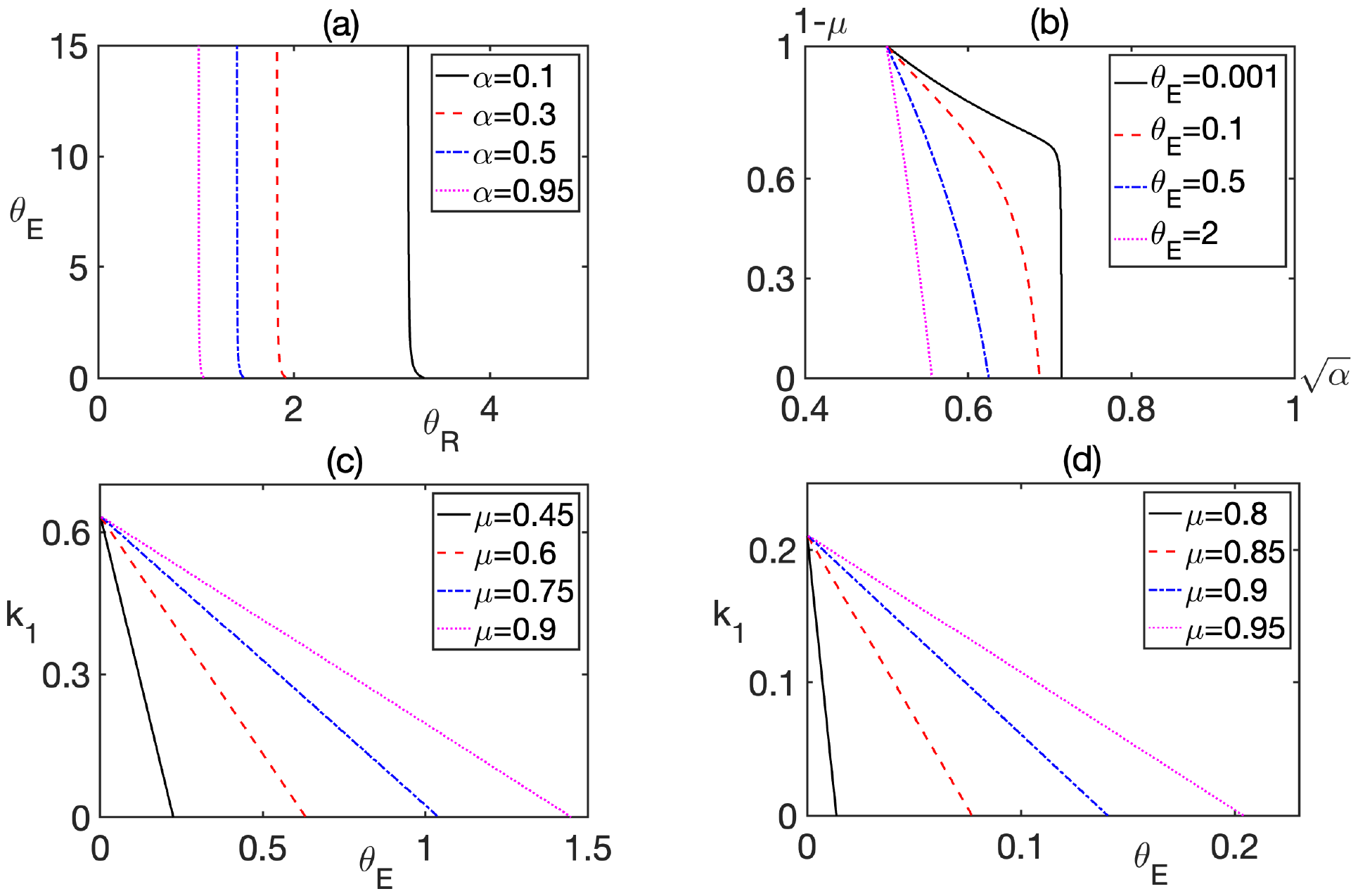
Boundaries between regions of stability of extinction steady state *E*_0_ and the mixed steady state *E*_2_. In all cases, the extinction steady state *E*_0_ is stable to the left of the boundary, and *E*_2_ is stable to the right of the boundary. (a) *µ* = 0.05, *k*_1_ = 0.7. (b) *θ*_*R*_ = 2, *k*_1_ = 0.7. (c) *θ*_*R*_ = 5, *α* = 0.1 (close to SMR). (d) *θ*_*R*_ = 5, *α* = 0.9 (close to GR).

We have also investigated how transition to lethal mutagenesis changes with mutant fitness, as represented by the parameter *k*_1_, and the rate *θ*_*E*_ characterising ExoN efficiency. Qualitatively, the picture is similar both in SMR, and in GR regimes (plots 3(c) and (d)), with higher rates of mutation *µ* resulting in larger parts of parameter plane, where the extinction steady state is stable. The notable difference between SMR and GR regimes is in that for the same values of all other parameters, the extinction steady state in the GR regimes is only observed for significantly smaller values of *θ*_*E*_, as well as smaller values of *k*_1_. A possible explanation for this observation is that since in the GR replication, both positive and negative RNA strands are used for subsequent viral replication, when ExoN is able to successfully correct replication errors in both of the strands, this proves quite effective at recovering the master sequence and subsequent replication, thus maintaining the mixed steady state *E*_2_. We also note that as mutations become more neutral, i.e. the value of *k*_1_ increases, differences between scenarios with different mutation rates become less notable.

## 4. Modelling the effects of treatment on viral replication

Over the years, various strategies have been proposed to minimise epidemiological burden of diseases caused by CoV, as well as to prevent severe disease and death in infected individuals. While vaccines, both vector-based and, most recently, mRNA, provide protection by means of eliciting virus-specific T cells and antibodies that are able to destroy infected cells and kill free virions, clinical treatments have largely focused on alleviating symptoms and/or reducing within-patient viral replication. Despite significant progress in treatment of diseases cause by different RNA viruses, there have not yet been drugs approved that were developed specifically to treat disease caused by CoV, though during a recent COVID-19 pandemic, a number of earlier developed drugs have been repurposed for treatment [69, 89], and most of the examples we consider below come from the search for successful antivirals against SARS-CoV-2. One promising antiviral target is the RdRp [115, 107, 112], an essential protein required for within-cell viral replication in RNA viruses, and a number of drugs have been shown to act as RdRp inhibitors. RdRp inhibition has already proved effective for hepatitis B and C [43, 17], as well as for HIV [72]. Among these drugs, a major class is the so-called *nucleoside analogues* (NAs) - these are pro-drugs that are metabolised/phosphorylated into an active 5’-triphosphate form, which is able to compete with endogenous nucleotides for being incorporated by the RdRp into the nascent viral RNA strand [89, 1]. Once this happens, subsequent synthesis of viral RNA chain is stopped either immediately (such drugs are called immediate or obligate chain terminators), or further down the growing RNA chain (delayed chain terminators) [108].

A major obstacle for successful action of NAs as RdRp inhibitors is their excision by CoV ExoN [89, 112, 109], and different NA drugs have different degree, at which they can be excised by the ExoN, with, for instance, sofosbuvir being more resistant to excision than remdesivir [47], and ribavirin being very easily excised [114, 34, 108], while tenofovir being largely resistant to excision [30, 114, 109]. Recent studies have identified some of the structural features of NAs that can make them more resistant to ExoN activity [109, 112, 10, 21]. It was shown that nucleotides possessing both 2’- and 3’-OH groups were more efficiently removed by the ExoN, while nucleotides lacking both of those groups were more resistant to excision [109]. Furthermore, the presence the 3’-OH group in the NA was more critical than the 2’-OH for excision by ExoN [109]. One strategy for achieving RdRp inhibition while avoiding excision by the ExoN is by using non-nucleoside RdRp inhibitors, majority of which bind to allosteric sites on the surface of RdRp, thus changing its conformation and hence, affecting its ability to bind to substrates [59, 95, 19].

Being RNA viruses, CoV are characterised by high mutation rates, and their replication fidelity is primarily achieved by the correcting action of ExoN. This suggests that ExoN itself can be considered to be a viable antiviral target [50, 66, 78]. So far, no drugs have been proposed as specific ExoN inhibitors. However, during the course of COVID-19 pandemic, several drugs have been identified as tentative ExoN inhibitors either through molecular docking simulations [50], or by observing clinical efficacy of using certain drugs in addition to known NAs that would otherwise be readily excised by the ExoN [108]. Deval and Gurard-Levin [21] provide a nice overview of recent studies that have extensively explored both known NAs and hundreds of small molecules to identify the most promising candidates for ExoN inhibitors. Very recently, some NA have been proposed to simultaneously have the dual action of both RdRp and ExoN inhibitors, which would make them very strong potential antiviral against CoV [109, 77]. Table 1 contains some examples of RdRp inhibitors, including NAs (both immediate and delayed chain terminators) and non-nucleoside inhibitors, as well as ExoN inhibitors.

**Table 1:** Single action RdRp and ExoN inhibitors.

| RdRp inhibitors |  |  |  |
| --- | --- | --- | --- |
| Nucleoside analogues |  |  |  |
| Obligate/Immediate | Non-obligate/Delayed | Non-nucleoside | ExoN inhibitors |
| Tenofovir[108], sofosbuvir[108], stavudine[109], zidovudine[109] | Remdesivir[89, 52, 38] entecavir[38] | Dasabuvir[114, 107], suramin[112] corilagin[112], lycorinne[112] | Pibrentasvir[108], ombitasvir[108] hesperidin[50], conivaptan[50] |

An alternative antiviral strategy that is particularly relevant for CoV is based on lethal mutagenesis, where instead of inhibiting viral replication by targeting RdRp, the idea is instead to further increase the rate of viral mutations during replication to overwhelm proof-reading and fidelity-inducing action of ExoN, thus causing lethal mutagenesis and extinction of the virus [73]. While lethal mutagenesis has already been extensively studied clinically for other RNA viruses [14, 56, 91, 70, 41, 79], applying this strategy to CoV has so far been quite limited [90, 48], and it has only been recently shown experimentally that coronaviruses with absent or ineffective ExoN activity are, in fact, susceptible to lethal mutagenesis [93]. One issue that affects the feasibility of this strategy for CoV is the same as with other NAs - these drugs are themselves quite well excised by ExoN, and hence, the success of lethal mutagenesis significantly depends on giving the drugs the time to act by suppressing proofreading action of ExoN [90, 48]. Among known NAs, so far only ribavirin has been suggested to act through both RdRp inhibition, and by causing lethal mutagenesis [14, 89], though it is also known to be effectively excised by the ExoN [69, 34]. We have collected in Table 2 examples of mutagenic NAs, as well dual inhibitors of RdRp and ExoN, and RdRp inhibitor with lethal mutagenesis.

**Table 2:** Mutagenic NAs and combined action drugs.

| Mutagenic NAs | Combined RdRp and ExoN inhibitors | Combined RdRp inhibition and lethal mutagenesis |
| --- | --- | --- |
| Favipiravir[108, 112], molnupiravir[114, 112, 48] | Velpatasvir[109], daclatasvir[109] riboprine[77], forodesine[77] | Ribavirin[14, 89] |

Mathematically, the effects of all these different types of drugs can be represented in our model through their effect on RdRp and ExoN efficacy, as represented by parameters *θ*_*R*_ and *θ*_*E*_, respectively, for RdRp inhibitors and ExoN inhibitors, or on mutation rate *µ* for mutagenic drugs. Importantly, as already mentioned, since many of the drugs are NAs, they themselves are to some degree susceptible to excision by ExoN. We model the excision of various NA drugs by ExoN by introducing a factor

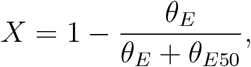

which quantifies the fraction of drug that is not excised, depending on half-maximum value *θ*_*E*50_. As *θ*_*E*50_ approaches the values or 0 or *∞*, we obtain, respectively, regimes, where drugs are fully excised or not excised at all. The latter situation, represented by *X* = 1, also applies to the case of non-nucleoside RdRp inhibitors.

Effects of RdRp inhibitors (RI) can be represented in the model by replacing the rate of *θ*_*R*_

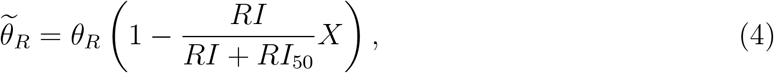

with where the second term represents the effect of the inhibitor given its concentration *RI* and half-maximum concentration *RI*_50_, subject to its possible excision by ExoN. Obviously, increasing the concentration of RI drugs (or, equivalently, reducing *RI*_50_) increases the level of RdRp inhibition. Similarly, pure ExoN inhibitors (EI) act by suppressing excision activity, which results in replacing *θ*_*E*_ by

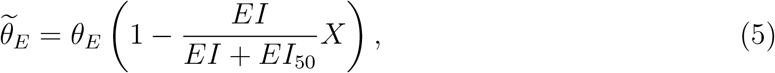

with a similar analogy in that higher EI drug concentrations (or lower *EI*_50_) leads to a higher level of ExoN excision activity. For both RI and EI drugs, we account for the fact that they can possibly be excised by ExoN, which is modelled by including the rate *X* that quantifies how much of the drug remains non-excised.

In terms of possible interactions between RI and EI drugs, several clinical protocols have been implemented that demonstrated how efficacy of RI drugs could be enhanced if patients were also simultaneously given some EI drugs [108]. Furthermore, the drugs with combined RI and EI action have been screened [109, 77], and their combined effect consists in replacing both *θ*_*R*_ and *θ*_*E*_ by, respectively, 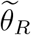 and 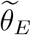 with the same drug concentration, i.e. *RI* = *EI*. However, since RI and EI drugs may be excised by ExoN at different rates, this will be accounted for by writing

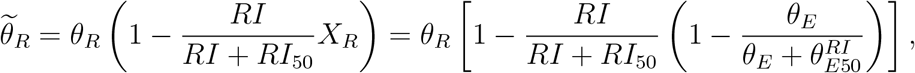

and

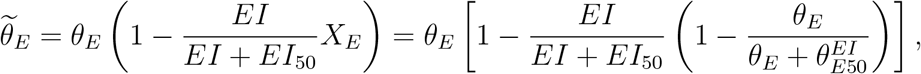

where 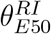 and 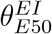 are half-maximal effective rates of excising RI and EI drugs, respectively.

To model the effects of NAs on viral replication dynamics, we separately consider two different types of NA-induced effects. The first one works by means of reducing the rate of strand replication due to chain termination. In the model, this would be represented by multiplying *θ*_*R*_ by a factor

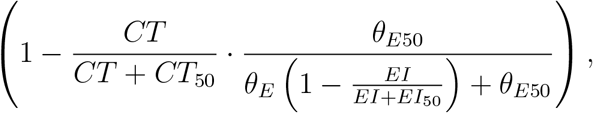

where *CT* and *EI* denote concentrations of chain-terminating and ExoN inhibiting drugs, respectively, *CT*_50_ and *EI*_50_ represent their respective half-maximal effective concentrations, and similarly, *θ*_*E*50_ is the half-maximum concentration of ExoN *θ*_*E*_, when it is acting as a inhibitor of CT through excising incorrect nucleosides incorporated into the growing RNA chain.

Another possibility is when drugs act by means of increasing mutation rate during strand replication, thus potentially leading to lethal mutagenesis. In the model, this can be represented by replacing the mutation rate *µ* by

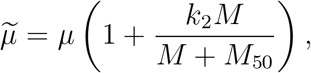

where *M* is the concentration of mutagenic drug, and *MG*_50_ is its half-maximal effective concentration. There is a natural restriction on the value of parameter *k*_2_, as given by 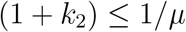 to ensure that the increased rate of mutation does not exceed 1.

Figure 4 illustrates critical curves separating regimes of presence of viral QS (stable steady state *E*_2_) and regimes of viral extinction (stable steady state *E*_0_) for different types of single- and dual action RdRp and ExoN inhibitors. For pure NA-based RdRp inhibitors, for each given value of ExoN efficacy *θ*_*E*_, there is some minimum half-maximum concentration *θ*_*E*50_, below which the drug is excised irrespective of its half-maximum concentration, and hence, the viral population persists. This minimum half-maximum concentration *θ*_*E*50_ itself increases with ExoN efficacy *θ*_*E*_, because higher *θ*_*E*_ indicates stronger excising efficacy of ExoN. For pure NA-based RdRp inhibitors, such as remdesivir, tenofovir, sofosbuvir etc., increasing *θ*_*E*_ results in reducing the half-maximum concentration *θ*_*E*50_ associated with a transition from the sustained viral state to viral extinction, as illustrated in Fig. 4(a), meaning that ExoN is becoming more effective at excising these drugs. Exactly the same picture is observed in Fig. 4(c) for pure ExoN inhibitors, such as pibrentasvir, ombitasvir and hesperidin. In the case of non-nucleoside RdRp inhibitors (which formally correspond to the case *θ*_*E*50_*→ ∞*), such as dasabuvir, suramin, corilagin, these drugs are not excised by ExoN at all, and hence, the excision factor is *X* = 1. Figure 4(b) shows that increasing the concentration *RI* of these drugs allows for a larger half-maximum concentration *RI*_50_, at which viral population can be eliminated through inhibition of RdRp. Finally, when a combination of RdRp and ExoN inhibitors is used, as presented in Fig. 4(d), we observe a balance between half-maximal effective rates of excision of RI and EI drugs, where a higher half-maximum rate 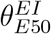 of excising ExoN inhibitors is associated with a lower half-maximum 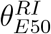 of excising RdRp inhibitors, at which a transition from the viral QS to extinction takes place. Making RdRp more effective at strand replication, i.e. increasing *θ*_*R*_, means that excision of RI drugs has to be more effective 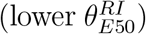 for the same rate of excision of EI drugs. In the case, where RdRp inhibition and ExoN inhibition is achieved by the same drugs, some examples of which are mentioned in Table 2, one can use *RI* = *EI* to represent the equality of concentrations of these drugs, which is the case we used for illustration in Fig. 4(d).

**Figure 4.**
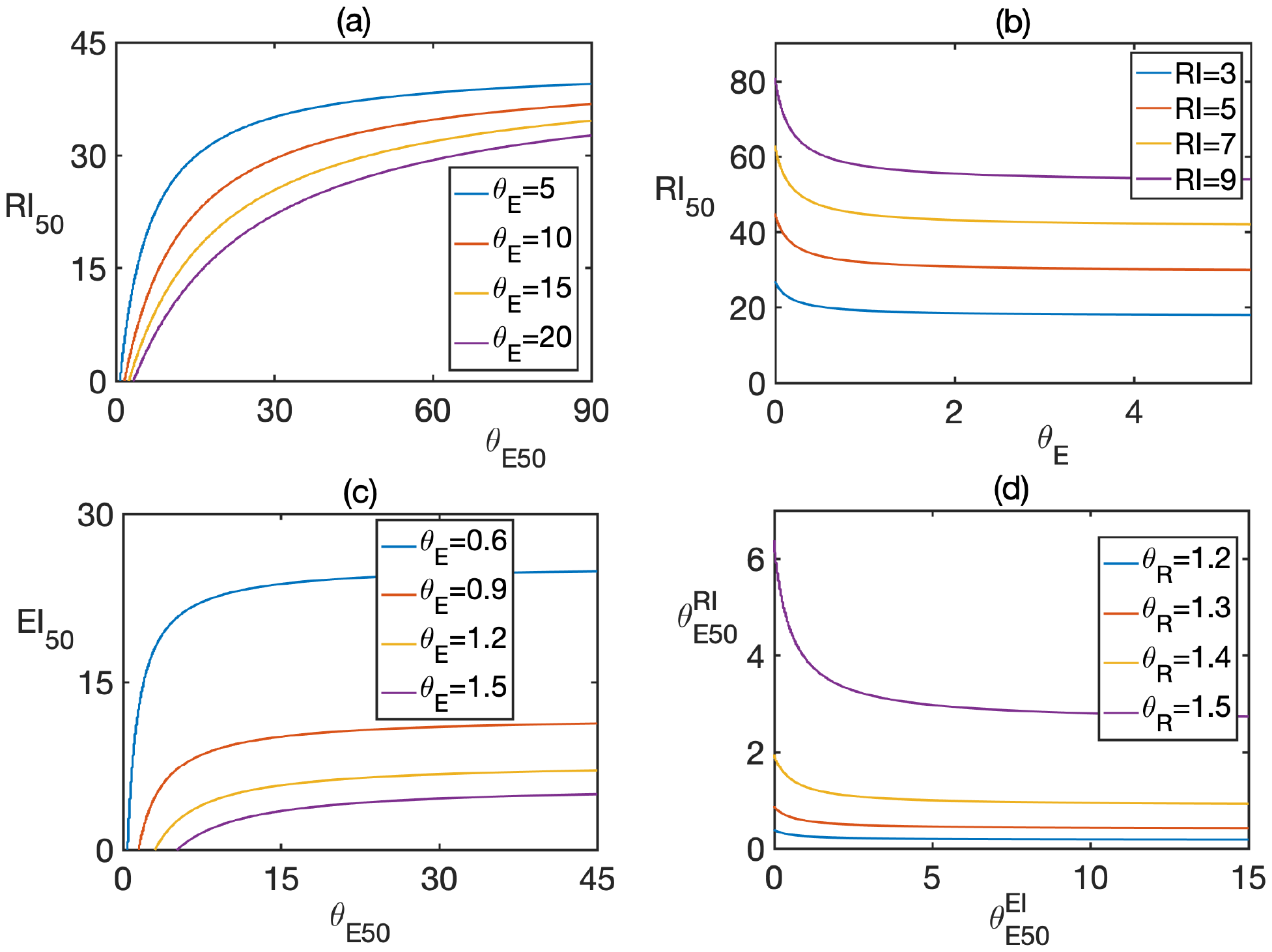
(a) Treatment with NA RdRp inhibitors. (b) Treatment with non-nucleoside RdRp inhibitors. (c) Treatment with pure ExoN inhibitors. (d) Treatment with a combination of RdRp and ExoN inhibitors. The steady state *E*_0_ with both strands being extinct is stable below the curves in plots (a)-(c) and above the curves in plot (d). Parameter values are *µ* = 0.05, *α* = 0.95, *k*_1_ = 0.7. (a) *θ*_*R*_ = 1.2, *RI* = 7. (b) *θ*_*R*_ = 1.2. (c) *θ*_*R*_ = 1.05, *EI* = 20. (d) *θ*_*R*_ = 1.5, *θ*_*E*_ = 0.6, *EI* = 20, *EI*_50_ = 20, *RI* = 20, *RI*_50_ = 40.

With clinical data suggesting high antiviral efficacy of certain drugs acting through lethal mutagenesis, in Fig. 5 we explore the effects of mutagenic drugs that can also be excised by the ExoN (favipiravir and molnupiravir), as well as drugs that both increase the rates of viral mutations, and also inhibit RdRp, such as ribavirin. While mutagenic drugs increase the rate of mutations, naturally, it cannot exceed 1, which imposes the restriction on parameter *k*_2_, characterising multiplicative increase in mutation rate. We have chosen basic mutation rate to be *µ* = 0.05, in which case the condition 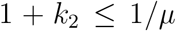 turns into *k*_2_ *≤* 19. For mutagenic drugs, as shown in Fig. 5(a), for each particular value of *k*_2_, there is a minimum half-maximum concentration *θ*_*E*50_, with mixed steady state *E*_2_ being stable for lower values of *θ*_*E*50_ for any half-maximum concentration of the drug. Biologically, this represents the case when the drug is efficiently excised irrespectively of its concentration before it is able to cause viral extinction through lethal mutagenesis. As with earlier discussion of other drugs, we observe a certain ‘tug-of-war’: increasing the rate *k*_2_, which controls how much the mutation rate is increased, leads to a reduction in the minimum value of *θ*_*E*50_, i.e. ExoN should be much more effective at excising highly mutagenic drugs in order to maintain the viral population and prevent it from going extinct. Figure 5(b) illustrates the treatment with a drug acting simultaneously through RdRp inhibition and lethal mutagenesis (such as ribavirin), but since this is one and the same drug, we use *RI* = *M* when computing the effective rates 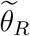and 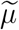.

**Figure 5.**
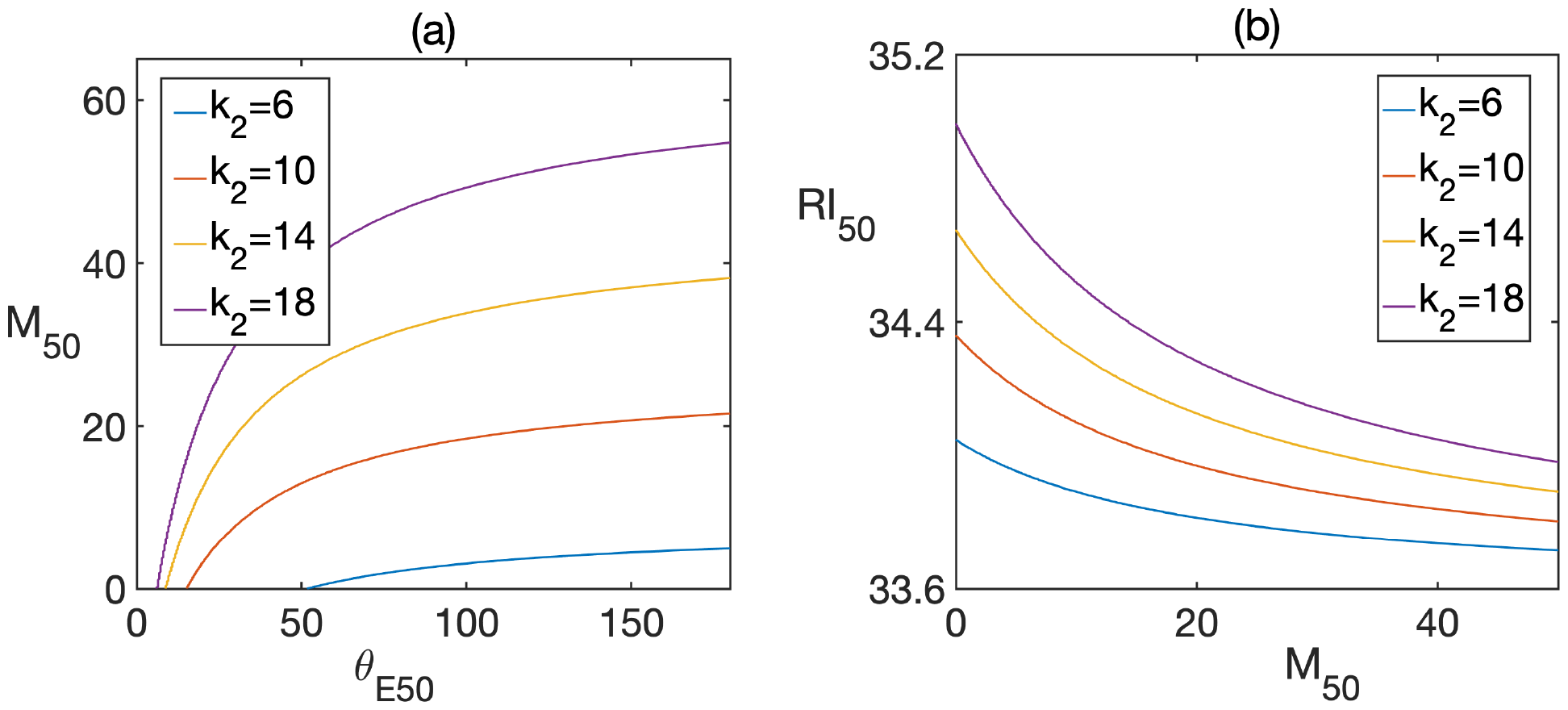
(a) Treatment with mutagenic drugs. (b) Treatment with a combination of mutagenic and ExoN-inhibiting drugs. Below the curves, the extinction steady state *E*_0_ is stable, above the curves the mixed steady state *E*_2_ is stable. Parameter values are *µ* = 0.05, *α* = 0.95, *k*_1_ = 0.7 (a) *θ*_*R*_ = 1.03, *θ*_*E*_ = 20, *M* = 20. (b) *θ*_*R*_ = 1.4, *θ*_*E*_ = 50 = 50, *RI* = 20.

## 5. Discussion

In this paper we have studied a mathematical model of within-cell replication of CoV with account for mutations and the role of exoribonuclease in improving replication fidelity through recognising and excising incorrect nucleotides attached to the growing RNA chains by error-prone RdRp. The majority of earlier theoretical work on replication dynamics of RNA viruses focused on understanding their evolutionary dynamics from the perspective of critical thresholds associated with error catastrophe and lethal mutagenesis. We recall the fundamental distinction between these two concepts: error catastrophe refers to an evolutionary change in the genomic space where master sequences are outcompeted by the mutants. In contrast, lethal mutagenesis refers to the idea where a large accumulation of mutants with low fitness leads to the extinction of the entire viral population. We have explored both of these scenarios for the case, where ExoN is either absent or non-functional, as well as when it does have an effect. Under an assumption of flat Swetina-Schuster fitness landscape, where all mutants are considered to have exactly the same fitness, increasing the mutation rate reduces the parameter region, where the viral quasispecies is able to survive and avoid extinction. Earlier studies showed that viral replication mode that controls which RNA strands are used for subsequent replication, can have a profound influence on viral persistence in terms of virus ability to withstand negative effects associated with deleterious mutations. Although these is some experimental evidence that certain viruses are characterised by a replication mode that is closer to SMR or GR mode, for the majority of RNA viruses, including CoV, actual replication mode is unknown, and it is quite likely that these viruses lie somewhere in the middle between these two replication modes. We have shown how viral quasispecies can disappear through either an error catastrophe or lethal mutagenesis for different replication modes, with higher chances of virus survival for the replication mode closer to GR. Importantly, whenever ExoN is functional, this prevents the mutant-only steady state associated with reorganisation of the genomic space from being feasible, as ExoN would be able to excise incorrect nucleotides from the mutant, thus recovering the master strand. Our results show that increasing the rate, at which ExoN is able to excise incorrect nucleotides from the growing RNA chain, results in making the viral quasispecies persistent for lower values of the parameter *α*, i.e. for replication modes closer to SMR than to GR. The closer the virus replication mode is to geometric replication, the larger is the range of values of the parameter *θ*_*R*_ that controls the replication strength of RdRp, for which viral quasispecies can persist for the same excising efficacy of ExoN.

With CoV being quite unique in having the largest known genomes among RNA viruses, and being known for having caused significant global pandemics over the last 20 years, it is essential to understand how antivirals can effectively reduce within-cell replication of CoV, based on the strategies of error catastrophe and lethal mutagenesis. With RdRp being en essential ingredient of viral replication, and ExoN playing a crucial error-correcting role, both of these enzymes have been suggested as potential targets for antivirals. Moreover, lethal mutagenesis has been shown to be an effective antiviral strategy for other RNA viruses, hence, it was natural to also explore its feasibility for CoV. We have analysed the effects of different types of antivirals, including RdRp inhibitors based on nucleoside analogues, non-nucleoside RdRp inhibitors, tentative ExoN inhibitors, as well as mutagenic drugs. In each case we showed how, depending on the balance between a particular drug’s mode of action (inhibition of RdRp or Exon, or increased mutation) and drug’s excision by the viral ExoN, it is either possible to achieve virus elimination through lethal mutagenesis, or the virus is able to withstand treatment due to a combination of mutations and efficient drug excision. The importance of these results lies in demonstrating the specific role played by ExoN in reducing the efficacy of antiviral drugs.

There is a number of interesting and exciting directions, in which the model studied in this paper could be extended to achieve better realism, as well as to provide a more comprehensive understanding of within-cell viral dynamics. One fundamental assumption of our model concerned the structure of fitness landscape, which for simplification was chosen to be a flat Swetina-Schuster landscape, which has been successfully used in a number of earlier studies to model the dynamics of RNA viruses [5, 71, 96]. This also fits with the ‘survival-of-the-flattest’ paradigm, in which quasispecies with high mutation rates survive through having the flattest possible fitness landscape that provides them with the highest degree of robustness against deleterious mutations [11, 103]. In many cases, however, the fitness landscape is much more complex [53, 84], and one could include into our model differential fitness of mutants with different numbers (and types) of mutations, and explore their possible cooperation/competition for cell resources. Some of the fitness landscapes that could be investigated in the context of our model, and for which the survival and possible extinction of viral species have been studied earlier, include the multiplicative landscape [6, 7], multivariate Gaussian landscape [62], and a biophysical landscape [9], in which the fitness landscape was determined experimentally through observation of how mutations affect thermal stability of proteins. Unlike the multiplicative landscape, the biophysical fitness landscape allows for both beneficial and deleterious mutations, and it was subsequently shown to agree quite well with the observation of mutational effects in RNA viruses [110]. Reducing the size of viral population with such fitness landscape was shown to reduce the rate of mutations that are required to make the viral population go extinct [111]. In the absence of robust data on fitness of CoV, one of the next steps could be an exploration of how these different fitness landscapes would affect the dynamics of CoV replication.

In order to better understand CoV dynamics, it is important to pinpoint precise mechanism, through which lethal mutagenesis can result in CoV losing its viability and becoming extinguished. While conventional representation of the error catastrophe is based on the idea where genome viability is lost once the accumulated number of errors exceeds certain error threshold, experimental results suggest that the relation between mutagen exposure and virus viability is often significantly more complex [100]. More specifically, genome inactivation is often, if not predominantly, the result of a single lethal mutation rather than an accumulation of multiple deleterious mutations [37]. For example, in the vesicular stomatitis virus, only 30% of mutations were deleterious (and not lethal), and they reduced viral growth rate by, on average, 19%, while almost 40% of random single nucleotide substitutions were actually lethal [81], thus suggesting that virus viability may be much more strongly affected by single-hit lethal genome errors rather than by a sequential reduction in viral growth rates. This could be included in our model by considering separate subpopulations of mutants with different types of mutations, which is related to the above-mentioned problem of non-flat fitness landscapes.

Besides focusing on purely deleterious or lethal mutations, another mechanisms of how lethal mutagenesis can result in suppressing the virus is the so-called *lethal defection* [63, 39, 40, 67, 103, 104]. The idea of lethal defection is that rather than directly cause extinction of viral population by targeting its replicative capacity, hence reducing overall RNA virus population, mutagenesis can instead substantially increase virus population but create a large subpopulation of mutants, knows as defectors, that helps extinguish the virus through enhanced mutagenesis caused by these residually replication-competent mutants [63], as was shown for the first time experimentally in LCMV [39, 40]. This work built on an earlier observation of a complex nonlinear relation between mutational load and extinction of viral population [41, 91]. Iranzo and Manrubia [45] developed a stochastic model of lethal defection and showed how for small virus population sizes, even if mutations rates themselves are low, once defectors are present, their fixation in population due to genetic drift eventually results in the displacement of viable virus population, thus causing extinction of the virus. While providing a feasible mechanism of stochastic virus extinction, this model only applies when the size of viral population is relatively small, which is known not to be the case for some of real viruses [39, 40]. Moreover, this model assumes that mutations affect virus replicative capacity and infectivity in a synchronised manner, which may also not always be the case. Moreno et al. [67] looked into whether lethal defection based on interference-complementation interactions between defectors and the rest of the virus is able to explain how initial multiplicity of infection can result in lethal mutagenesis. Importantly, the degree of interference is known to be affected by the virus replication mode [84, 87], and our model could be modified to study lethal defection with account for virus replication mode.

An important practical issue that can be studied within the framework of our model is the application of mutagenic drugs at sublethal doses, as well as more complex drug interactions. While aiming to reduce burden on patients, applying mutagenic drugs at sublethal doses can actually prove to be counter-productive from a clinical perspective, as it may help the virus to escape immune response. This can result in the emergence of mutants that would be resistant to a variety of non-mutagenic and mutagenic drugs, as has been known to happen with antibiotic-resistant bacteria [75]. Rather than being merely a theoretical consideration, this effect has already demonstrated in ribavirin, a well-known broad spectrum antiviral, when used for the treatment of FMDV. Experiments show that when ribavirin was given to patients at high concentrations from the very start of treatment, this led to a successful virus elimination [74]. In contrast, providing ribavirin in a sequence of increasing concentrations, led to the emergence of ribavirin-resistant mutants [92]. In all of the simulations presented in this paper, whenever we considered the simultaneous use of mutagenic and replication-inhibiting drugs, these were assumed to be used at the same time. The problem is that this can actually be a detrimental approach, as at certain dosages, these drugs are antagonistic - by increasing the rate of mutations, mutagenic drugs can make mutants more resistant to inhibitors [46]. As an alternative, it was, therefore, suggested to first try to use replication inhibitors, and only then use mutagens, to achieve maximum efficacy of virus elimination [74, 61, 46]. Our model could be used for testing the effects of sublethal dosages of mutagenic drugs, and also for analysing and identifying optimal treatment regimens for combinations of mutagenic and replication inhibiting drugs when used for treatment of CoV infections.

Finally, the model presented in this paper focused on within-cell replication-mutation dynamics and used a mean-field deterministic representation of viral species dynamics. With mutations being intrinsically random, over the years a number of models have been proposed that tried to capture stochastic nature of virus replication dynamics with mutations. This included, among others, a stochastic branching model of Demestrius [18] that was subsequently extended into a phenotypic model [3, 31]. Dalmau [16] proposed an alternative stochastic model of quasispecies based on multi-type branching processes. Sardanyés et al.[86] used a stochastic model to explore the effects of various fitness landscapes on virus survivability under different types of mutations, demonstrating how variable distributions of mutational fitness effects may prevent or reduce the chances of viral extinction by providing sufficient amount of replication-capable phenotypes. Loverdo and Lloyd-Smith [57] also studied survival and invasion of lineages with beneficial or deleterious mutations. Since viruses engage in complex interactions with the immune system of their hosts, one could also look at higher-level models that include not only intracellular, but also extracellular virus dynamics and different types of immune response. A number of stochastic models have looked at this problem [54, 113, 32], but those models rather focused on studying stochastic aspects of immune responses, and not accounting for changes in virus population due to mutations, and the effects of mutations on virus replication. In this respect, one potential direction of future research is the development of a stochastic version of our model that would allow one to explore in more detail the role of small virus population sizes, to get a better idea of the distributions of potential genomic profiles of the viral quasispecies under mutations and different fitness landscapes, and to analyse interactions between the virus and the immune system. This is particularly important from the perspective of understanding and developing better antiviral treatments, as lethal mutagenesis is largely associated with stochastic extinction occurring at low population sizes that result from accumulation of low-fitness mutants.

## References

[1] Agostini, M.L, Andres, E.L., Sims, A.C. et al., 2018. Coronavirus susceptibility to the antiviral remdesivir (GS-5734) is mediated by the viral polymerase and the proofreading exoribonuclease. mBio 9: e00221–18.

[2] Alonso, J., Fort, H., 2010. Error catastrophe for viruses infecting cells: analysis of the phase transition in terms of error classes. Phil. Trans. R. Soc. A 368, 5569–5582.

[3] Antoneli, F., Bosco, F.A.R., Castro, D., Janini, L.M.R., 2013. Virus replication as a phenotypic version of polynucleotide evolution. Bull. Math. Biol. 75, 602–628.

[4] Binder, M., Sulaimanov, N., Clausznitzer, D., Schulze, M., Hüber, C.M. et al., 2013. Replication vesicles are load- and choke-points in the hepatitis C virus lifecycle. PLoS Pathog 9: e1003561.

[5] Bull, J.J., Meyers, L.A., Lachmann, M., 2005. Quasispecies made simple. PLoS Comput. Biol. 1, e61.

[6] Bull, J.J., Sanjuán, R., Wilke, C.O., 2007. Theory of lethal mutagenesis for viruses. J. Virol. 81, 2930–2939.

[7] Bull, J.J, Sanjuán, R., Wilke, C.O. 2008. Lethal mutagenesis. In: Domingo, E., Parrish, C.R., Holland, J.J. (eds) Origin and evolution of viruses. Elsevier Academic Press, Amsterdam.

[8] Chao, L., Rang, C.U., Wong, L.E., 2002. Distribution of spontaneous mutants and inferences about the replication mode of the RNA bacteriophage ϕ6. J. Virol. 76, 3276–3281.

[9] Chen, P, Shakhnovich, E., 2009 Lethal mutagenesis in viruses and bacteria. Genetics 183, 639–650.

[10] Chinthapatla, R., Sotoudegan, M., Srivastava, P., 2023. Interfering with nucleotide excision by the coronavirus 3’-to-5’exoribonuclease. Nucl. Acids Res. 51, 315–336,

[11] Codoñer, F.M., Daros, J.-A., Solé, R.V., Elena, S.F., 2006. The fittest versus the flattest: experimental confirmation of the quasispecies effect with subviral pathogens. PLoS Pathog. 2, e136.

[12] Combe, M., Garijo, R., Geller, R., Cuevas, J.M., Sanjuán, R., 2018. Single-cell analysis of RNA virus infection identifies multiple genetically diverse viral genomes within single infectious units. Cell Host Micr. 18, 424–432.

[13] Cornish-Bowden, A., 2015. One hundred years of Michaelis-Menten kinetics. Persp. Sci. 4, 3–9.

[14] Crotty, S., Cameron, C.E., Andino, R. (2001), RNA virus error catastrophe: direct molecular test by using ribavirin, Proc. Natl. Acad. Sci. U.S.A. 98, 6895–6900.

[15] Dahari, H., Ribeiro, R.M., Rice, C.M., Perelson, A.S., 2007, Mathematical modeling of subgenomic hepatitis C virus replication in Huh-7 cells. J. Virol. 81, 750–760.

[16] Dalmau, J., 2016, Distribution of the quasispecies for a Galton-Watson process on the sharp peak landscape. J Appl. Probab. 53, 606–613.

[17] Das, D., Pandya, M., 2018. Recent advancement of direct-acting antiviral agents (DAAs) in Hepatitis C Therapy. Mini-Rev. Med. Chem. 18, 584–596.

[18] Demetrius, L., 1985. The units of selection and measures of fitness. Proc. R. Soc. Lond. B 225, 147–159.

[19] Dejmek, M., Konkolová, E., Eyer, L. et al., 2021. Non-nucleotide RNA-dependent RNA polymerase inhibitor that blocks SARS-CoV-2 replication. Viruses 13, 1585.

[20] Denison, M.R., Graham, R.L., Donaldson, E.F., Eckerle, L.D., Baric, R.S., 2011. Coronaviruses, RNA Biol. 8, 270–279.

[21] Deval, J., Gurard-Levin, Z.A., 2022. Opportunities and challenges in targeting the proofreading activity of SARS-CoV-2 polymerase complex. Molecules 27, 2918.

[22] Domingo, E., Sabo, D., Taniguchi, T., Weissman, C. 1978. Nucleotide sequence heterogeneity of an RNA phage population. Cell 13, 735–744.

[23] Domingo, E., Holland, J., Ahlquist, P., 1988 RNA genetics. Boca Raton, FL, CRC Press.

[24] Domingo, E.,2000. Viruses at the edge of adaptation. Virol. 270, 251–253.

[25] Domingo, E., 2005. Antiviral strategy on the horizon. Virus Res. 107, 115–116.

[26] Domingo, E., Holland, J.J., 1994. Mutation Rates and Rapid Evolution of RNA Viruses. In: Morse, S.S. (ed.), The Evolutionary Biology of Viruses, Raven Press, New York, pp. 161–184.

[27] Drake, J.W., Holland, J.J., 1999. Mutation rate among RNA viruses. Proc. Natl. Acad. Sci. USA 96, 13910–13913.

[28] Eigen, M., 1971. Self-organization of matter and the evolution of biological macromolecules. Naturwiss 58, 465–523.

[29] Eigen, M., Biebricher, C.K., 1998. Sequence space and quasispecies distribution. In Domingo, E., Holland, J., Ahlquist, P., RNA genetics, Boca Raton, FL, CRC Press, pp. 211–245.

[30] Elfiky, A.A., 2020. Ribavirin, Remdesivir, Sofosbuvir, Galidesivir, and Tenofovir against SARS-CoV-2 RNA dependent RNA polymerase (RdRp): A molecular docking study. Life Sci. 253, 117592.

[31] Fabreti, L.G., Castro, D., Gorzoni, B., Janin, L.M.R., Antoneli, F., 2019. Stochastic modeling and simulation of viral evolution. Bull. Math. Biol. 81, 1031–1069.

[32] Fatehi, F., Kyrychko, S.N., Ross, A., Kyrychko, Y.N., Blyuss, K.B., 2018. Stochastic effects in autoimmune dynamics. Front. Physiol. 9, 45.

[33] Fehr, A.R., Perlman, S., 2015. Coronaviruses: an overview of their replication and pathogenesis. Coronaviruses 1282, 1–23.

[34] F. Ferron, L. Subissi, A. T. S. De Morais, Structural and molecular basis of mismatch correction and ribavirin excision from coronavirus RNA, Proc. Natl. Acad. Sci. USA 2017 115 (2) E162–E171.

[35] J. Fornés, J.T. Lázaro, T. Alarcon, S.F. Elena, Viral replication modes in single-peak fitness landscapes: A dynamical systems analysis, J. Theor. Biol. 460, 170–183 (2019)

[36] Garcia-Villada, L., Drake, J.W., 2012. The three faces of riboviral spontaneous mutation: spectrum, mode of genome replication, and mutation rate. PLoS Genet. 8, e1002832.

[37] Gard, S., Maaløe, O., 1959. Inactivation of viruses. In Burnet, M.F., Stanley, W.M. (ed.), The viruses. Academic Press, Orlando, Fl, p. 359–427.

[38] Gordon, C.J., Lee, H.W., Thesnokov, E.P. et al., 2022. Efficient incorporation and template-dependent polymerase inhibition are major determinants for the broad-spectrum antiviral activity of remdesivir. J. Biol. Chem. 298, 101529.

[39] Grande-Pérez, A., Gomez-Mariano, G., Lowenstein, P.R., Domingo, E., 2005. Mutagenesis-induced, large fitness variations with an invariant arenavirus consensus genomic nucleotide sequence. J. Virol. 79, 10451–10459.

[40] Grande-Pérez, A., Lazaro, E., Lowenstein, P., Domingo, E., Manrubia, S.C., 2005. Suppression of viral infectivity through lethal defection. Proc. Natl. Acad. Sci. USA 102, 4448–4452.

[41] Grande-Pérez, A., Sierra, S., Castro, M.G., Domingo, E., Lowenstein, P.R., 2002. Molecular indetermination in the transition to error catastrophe: systematic elimination of lymphocytic choriomeningitis virus through mutagenesis does not correlate linearly with large increases in mutant spectrum complexity. Proc. Natl. Acad. Sci. USA 99, 12938–12943.

[42] Grebennikov, D., Kholodareva, E., Sazonov, I., Karsonova, A., Meyerhans, A., Bocharov, G., 2021. Intracellular life cycle kinetics of SARS-CoV-2 predicted using mathematical modelling, Viruses 13, 1735.

[43] He, Z., Wang, J., Liu, K., Huang, H., Du, Y., Lin, Z. et al. 2012. Randomized trial of lamivudine, adefovir, and the combination in HBeAg-positive chronic hepatitis B. Gastroenterol. 36, 592–597

[44] Holland, J.J., Domingo, E., de la Torre, J.C., Steinhauer, D.A.,1990. Mutation frequencies at defined single codon sites in vesicular stomatitis virus and polio virus can be increased only slightly by chemical mutagenesis. J. Virol. 64, 3960–3962.

[45] Iranzo, J., Manrubia, S.C., 2009. Stochastic extinction of viral infectivity through the action of defectors. Eur. Phys. Lett. 85, 18001.

[46] Iranzo, J., Perales, C., Domingo, E., Manrubia, S.C., 2011. Tempo and mode of inhibitor-mutagen antiviral therapies: a multidisciplinary approach. Proc Natl Acad Sci USA 108, 16008–16013.

[47] Jockusch, S., Tao, C., Li, X. et al., 2020. Sofosbuvir terminated RNA is more resistant to SARSCoV2 proofreader than RNA terminated by Remdesivir. Sci. Reps. (2020) 10:16577

[48] Kabinger, F., Stiller, C., Schmitová J. et al., 2021. Mechanism of molnupiravir-induced SARS-CoV-2 mutagenesis. Nature Struct. & Mol. Biol. 28, 740–746.

[49] Kesheh, M.M., Hosseini, P., Soltani, S., Zandi, M., 2022. An overview on the seven pathogenic human coronaviruses. Rev. Med. Virol. 32, e2282.

[50] Khater, S., Kumar, P., Dasgupta N. et al., 2021 Combining SARS-CoV-2 proofreading exonuclease and RNA-Dependent RNA Polymerase inhibitors as a strategy to combat COVID-19: a high-throughput in silico screening. Front. Microbiol. 12, 647693.

[51] Kim, J.K., Tyson, J.J., 2020. Misuse of the Michaelis-Menten rate law for protein interaction networks and its remedy. PLoS Comput. Biol. 16, e1008258.

[52] Kokic, G., Hillen, H.S., Tegunov, D. et al., 2021. Mechanism of SARS-CoV-2 polymerase stalling by remdesivir. Nature Comms. 12, 279.

[53] Lalic, J., Elena, S.F., 2015. The impact of high-order epistasis in the within-host fitness of a positivesense plant RNA virus. J. Evol. Biol. 28, 2236–2247.

[54] Lipniacki, T., Paszek, P., Brasier, A. R., Luxon, B.A., Kimmel, M. 2006. Stochastic regulation in early immune response. Biophys. J. 90, 725–742.

[55] Liu, D.X., Liang, J.Q., Fung, T.S., 2021. Human coronavirus-229E, -OC43, -NL63, and -HKU1 (Coronaviridae). Encycl. Virol. 2, 428–440.

[56] Loeb, L.A., Essigmann, J.M., Kazazi, F., Zhang, J., Rose, K.D., Mullins, J.I., 1999. Lethal mutagenesis of HIV with mutagenic nucleoside analogs. Proc. Natl Acad. Sci. USA 96, 1492–1497.

[57] Loverdo, C., Lloyd-Smith, J.O., 2013. Evolutionary invasion and escape in the presence of deleterious mutations. PLoS ONE 8, e68179.

[58] Luria, S.E., 1951. The frequency distribution of spontaneous bacteriophage mutants as evidence for the exponential rate of phage production. Cold Spring Harbor Symp. Quant. Biol. 16, 1505–1511.

[59] Ma, Y., Wu, L., Shaw, N., Gao, Y., Wang, J., Sun, Y. et al., 2015. Structural basis and functional analysis of the SARS coronavirus nsp14-nsp10 complex. Proc. Natl. Acad. Sci. USA 112, 9436–9441.

[60] Malone, B., Urakova, N., Snijder, E.J., Campbell, E.A., 2022. Structures and functions of coronavirus replication-transcription complexes and their relevance for SARS-CoV-2 drug design. Nature Rev. Mol. Cell. Biol. 23, 21–39.

[61] Manrubia, S.C., Domingo, E., Lázaro, E., 2010. Pathways to extinction: beyond the error threshold. Phil. Trans. R. Soc. Lond. B 365, 1943–1952.

[62] Martin, G., Gandon, S. 2010. Lethal mutagenesis and evolutionary epidemiology. Phil. Trans. R. Soc. Lond. B 365, 1953–1963.

[63] Martín, V., Abia, D., Domingo, E., Grande-Pérez, A., 2010. An interfering activity against lymphocytic choriomeningitis virus replication associated with enhanced mutagenesis. J. Gen. Virol. 91, 990–1003.

[64] Martíenez, F., Sardanyés, J., Elena, S.F., Daros, J.A., 2011. Dynamics of a plant RNA virus intracellular accumulation: stamping machine vs. geometric replication. Genetics 188, 637–646

[65] McLean, A.K., Luciani, F., Tanaka, M.M., 2010. Trade-offs in resource allocation in the intracellular life-cycle of hepatitis C virus. J. Theor. Biol. 267, 565–572.

[66] Moeller, N.H., Shi, K., Demir, Ö., Aihara, H., 2022. Structure and dynamics of SARS-CoV-2 proof-reading exoribonuclease ExoN. Proc. Natl. Acad. Sci. USA. 119, e2106379119.

[67] Moreno, H., Tejero, H., de la Torre, J.C., Domingo, E., Martín, V., 2012. Mutagenesis-mediated virus extinction: virus-dependent effect of viral load on sensitivity to lethal defection. PLoS ONE 7, e32550.

[68] Nakabayashi, J. 2012. A compartmentalization model of hepatitis C virus replication: an appropriate distribution of HCV RNA for the effective replication. J. Theor. Biol. 300, 110–117.

[69] Ogando, N.S., Ferron, F., Decroly, E., Canard, B., Posthuma, C.C., Snijder, E.J., 2019. The curious case of the Nidovirus exoribonuclease: its role in RNA synthesis and replication fidelity. Front. Microbiol. 10, 1813.

[70] Pariente, N., Sierra, S., Airaksinen, A., 2005. Action of mutagenic agents and antiviral inhibitors on foot-and-mouth disease virus. Virus Res. 107 (2), 183–193.

[71] Pastor-Satorras, R., Solé, R.V., 2001. Field theory of a reactiondiffusion model of quasispecies dynamics. Phys. Rev. E 64, 051909.

[72] Pedersen, O.S., Pedersen, E.B., 1999. Non-nucleoside reverse transcriptase inhibitors: the NNRTI boom. Antivir. Chem. Chemother. 10, 285–314.

[73] Perales, C., Domingo, E., 2016. Antiviral strategies based on lethal mutagenesis and error threshold. Curr. Top. Microbiol. Immunol. 392, 323–339.

[74] Perales, C., Agudo, R., Tejero, H., Manrubia, S.C., Domingo, E., 2009. Potential benefits of sequential inhibitor-mutagen treatments of RNA virus infections. PLoS Pathog. 5, e1000658.

[75] Pillai, S., Wong, J., Barbour, J., 2008. Turning up the volume on mutational pressure: Is more of a good thing always better? (A case study of HIV-1 Vif and APOBEC3). Retrovirol. 5, 26.

[76] Pizzato, M., Baraldi, C., Sopetto G.B. et al., 2001. SARS-CoV-2 and the host cell: a tale of interactions. Front. Virol. 1, 815388.

[77] Rabie, A.M., Abdalla, M., 2023. Evaluation of a series of nucleoside analogs as effective anticoronaviral-2 drugs against the Omicron-B.1.1.529/ BA.2 subvariant: a repurposing research study. Medic. Chem. Res. 32, 326–341.

[78] Rona, G., Zeke, A., Miwatani-Minter, B. et al., 2022. The NSP14/NSP10 RNA repair complex as a Pan-coronavirus therapeutic target. Cell Death Differ. 29, 285–292.

[79] Ruiz-Jarabo, C.M., Ly, C., Domingo, E., de la Torre, J.C., 2003. Lethal mutagenesis of the prototypic arenavirus lymphocytic choriomeningitis virus (LCMV). Virology 308, 37–47.

[80] Saakian, D. B., Biebricher, C. K., Hu, C.-K. 2009 Phase diagram for the eigen quasispecies theory with a truncated fitness landscape. Phys. Rev. E 79, 041905

[81] Sanjuán, R., Moya, A., Elena, S.F., 2004. The distribution of fitness effects caused by single-nucleotide substitutions in an RNA virus. Proc. Natl. Acad. Sci. U.S.A. 101, 8396–8401.

[82] Sanjuán, R., Nebot, M.R., Chirico, N., Mansky, L.M., Belshaw, R., 2010. Viral mutation rates. J. Virol. 84, 9733–9748.

[83] Sanjuán, R., Domingo-Calap, P., 2016. Mechanisms of viral mutation. Cell Mol. Life Sci. 73, 4433–3338.

[84] J. Sardanyés, R.V. Solé, S.F. Elena, Replication Mode and Landscape Topology Differentially Affect RNA Virus Mutational Load and Robustness, J. Virol. 83, 12579–12589 (2009).

[85] J. Sardanyés, F. Martínez, J.-A. Daros, S.F. Elena, Dynamics of alternative modes of RNA replication for positive-sense RNA viruses, J. Roy. Soc. Interface 9, 768–776 (2012)

[86] Sardanyés, J., Simo, C., Marténez, R. et al. Variability in mutational fitness effects prevents full lethal transitions in large quasispecies populations. Sci Rep 4, 4625 (2014).

[87] Sardanyés, J., Elena, S.F., 2011. Quasispecies spatial models for RNA viruses with different replication modes and infection strategies. PLoS ONE 6 (9), e24884.

[88] Schulte, M.B., Draghi, J.A., Plotkin, J.B., Andino, R., 2015. Experimentally guided models reveal replication principles that shape the mutation distribution of RNA viruses. eLife 4, e03753.

[89] Shannon, A., Le, N. T.-T., Selisko B. et al., 2020. Remdesivir and SARS-CoV-2: structural requirements at both nsp12 RdRp and nsp14 exonuclease active-sites. Antivir. Res. 178, 104793.

[90] Shannon, A., Selisko, B., Le, NTT. et al. 2020. Rapid incorporation of favipiravir by the fast and permissive viral RNA polymerase complex results in SARS-CoV-2 lethal mutagenesis. Nat. Commun. 11, 4682.

[91] Sierra, S., Davila, M., Lowenstein, P.R., Domingo, E., 2000. Response of foot-and-mouth disease virus to increased mutagenesis: influence of viral load and fitness in loss of infectivity. J. Virol. 74, 8316–8323.

[92] Sierra, M., Airaksinen, A., Gonzalez-Lopez, C., Agudo, R., Arias, A., Domingo, E., 2007. Foot-and-mouth disease virus mutant with decreased sensitivity to ribavirin: implications for error catastrophe. J. Virol. 81, 2012–2024.

[93] Smith, E.C., Blanc, H., Vignuzzi, M., Denison, M.R., 2013. Coronaviruses lacking exoribonuclease activity are susceptible to lethal mutagenesis: evidence for proofreading and potential therapeutics. PLoS Pathog 9, e1003565.

[94] Smith, R.A., Loeb, L.A., Preston, B.D., 2005. Lethal mutagenesis of HIV. Virus Res. 107, 215–228.

[95] Sofia, M.J., Chang, W., Furman, P.A., Mosley, R.T., Ross, B.S., 2012. Nucleoside, nucleotide, and non-nucleoside inhibitors of hepatitis C virus NS5B RNA-dependent RNA-polymerase. J. Med. Chem. 55, 2481–2531.

[96] Solé, R.V., Sardanyés, J., Díez, J., Mas, A., 2006. Information catastrophe in RNA viruses through replication thresholds. J. Theor. Biol. 240, 353–359.

[97] Solé, R.V., Manrubia, S., Luque, B., Delgado, J., Bascompte, J., 1996. Phase transitions and complex systems. Complexity 1, 13–26.

[98] Solé, R.V., Goodwin, B.C., 2001. Signs of life: how complexity pervades biology. Basic Books, Perseus, New York.

[99] Stent, G., 1963. Molecular biology of bacterial Viruses. San Francisco, CA: W H Freeman and Company.

[100] Summers, J., Litwin, S., 2006. Examining the theory of error catastrophe. J. Virol. 80, 20–26.

[101] Swetina, J., Schuster, P., 1982. Self-replication with errors. A model for polynucleotide replication. Biophys. Chem. 16, 329–345.

[102] Tarazona, P., 1992. Error thresholds for molecular quasispecies as phase transitions: from simple landscapes to spinglass models. Phys. Rev. A 45, 6038–6049.

[103] Tejero, H., Marn, A., Montero, F., 2011. The relationship between the error catastrophe, survival of the flattest, and natural selection. BMC Evol. Biol. 11, 2.

[104] H. Tejero, F. Montero, J.C. Nuño, 2016.Theories of lethal mutagenesis: from error catastrophe to lethal defection. In Domingo, E., Schuster, P. (Eds.), Quasispecies: from theory to experimental systems. Springer International Publishing, Berlin, pp. 161–179.

[105] Thébaud, G., Chadouef, J., Morelli, M. J., McCauley, J. W., Haydon, D. T., 2010. The relationship between mutation frequency and replication strategy in positivesense single-stranded RNA viruses. Proc. R. Soc. B 277, 809–817.

[106] V’kovski, P., Kratzel, A., Steiner, S., Stalder, H., Thiel, V., 2021. Coronavirus biology and replication: implications for SARS-CoV-2, Nat. Rev. Microbiol. 19, 155–170.

[107] Uppal, T., Tuffo, K., Khaiboullina S. et al., 2022. Screening of SARS-CoV-2 antivirals through a cell-based RNA-dependent RNA polymerase (RdRp) reporter assay. Cell Insight 1, 100046.

[108] Wang, X., Sacramento, C.Q., Jockusch S. et al., 2022. Combination of antiviral drugs inhibits SARS-CoV-2 polymerase and exonuclease and demonstrates COVID-19 therapeutic potential in viral cell culture. Comms. Biol. 5, 154 (2022).

[109] Wang, X., Tao, C., Morozova I. et al., 2022. Identifying structural features of nucleotide analogues to overcome SARS-CoV-2 exonuclease activity. Viruses 14, 1413.

[110] Wylie, C.S., Shakhnovich, E.I., 2011. A biophysical protein folding model accounts for most mutational fitness effects in viruses. Proc. Natl. Acad. Sci. 108, 9916–9921.

[111] Wylie, C.S., Shakhnovich, E.I. 2012. Mutation induced extinction in finite populations: lethal mutagenesis and lethal isolation. PLoS Comput. Biol. 8, e1002609.

[112] Xu, X., Chen, Y., Lu X. et al., 2022. An update on inhibitors targeting RNA-dependent RNA polymerase for COVID-19 treatment: promises and challenges. Biochem. Pharmacol. 205, 115279.

[113] Yuan, Y., Allen, L.J., 2011. Stochastic models for virus and immune system dynamics. Math. Biosci. 234, 84–94.

[114] Zhao, J., Guo, S.S., Yi, D. et al., 2021. A cell-based assay to discover inhibitors of SARS-CoV-2 RNA dependent RNA polymerase. Antivir. Res. 190, 105078.

[115] Zhu, W., Chen, C.Z., Gorshkov, K., Xu, M., Lo, D.C., Zheng, W., 2020. RNA-Dependent RNA Polymerase as a target for COVID-19 drug discovery. SLAS Discov. 25, 1141–1151.

[116] Zitzmann, C., Schmid, B., Ruggieri, A., Perelson, A.S., Binder, M., Bartenschlager, R., Kaderali, L., 2020. A coupled mathematical model of the intracellular replication of dengue virus and the host cell immune response to infection. Front. Microbiol. 11, 725.

[117] Zitzmann, C., Kaderali, L., Perelson, A.S., 2020. Mathematical modeling of hepatitis C RNA replication, exosome secretion and virus release. PLoS Comput. Biol. 16, e1008421.

[118] C. Zitzmann, C. Dächert, B. Schmid et al., (2023). Mathematical modeling of plus-strand RNA virus replication to identify broad-spectrum antiviral treatment strategies. PLoS Comput. Biol. 19. e1010423.

